# Identification and comparison of individual chromosomes of three *Hordeum chilense* accessions, *Hordeum vulgare* and *Triticum aestivum* by FISH

**DOI:** 10.1101/255786

**Authors:** María-Dolores Rey, Graham Moore, Azahara C. Martín

## Abstract

Karyotypes of three accessions of *Hordeum chilense* (H1, H16 and H7), *Hordeum vulgare* and *Triticum aestivum* were characterized by physical mapping of several repetitive sequences. A total of fourteen repetitive sequences were used as probes for fluorescence in situ hybridization (FISH) with the aim of identifying inter‐ and intra-species polymorphisms. The (AG)_12_ and 4P6 probes only produced hybridization signals in wheat, the BAC7 probe only hybridized to the centromeric region of *H. vulgare*, and the pSc119.2 probe hybridized to both wheat and *H. chilense*, but not to *H. vulgare*. The remaining repetitive sequences used in this study produced a hybridization signal in all the genotypes. Probes pAs1, pTa535, pTa71, CCS1 and CRW were much conserved, showing no significant polymorphism among the genotypes studied. Probes GAA, (AAC)_5_, (CTA)_5_, HvT01 and pTa794 produced the most different hybridization pattern. We identified large polymorphisms in the three accessions of *H. chilense* studied, supporting the proposal of the existence of different groups inside *H. chilense* species. The set of probes described in this work allowed the identification of every single chromosome in all three species, providing a complete cytogenetic karyotype of *H. chilense, H. vulgare* and *T. aestivum* chromosomes, useful in wheat and tritordeum breeding programs.

## Introduction

*Hordeum chilense* Roem. et Schultz. (2n = 2× = 14; H^ch^ H^ch^) is a diploid wild barley, native of Chile and Argentina (von Bothmer et al. 1995), which shows very useful traits for wheat breeding, including drought and salt tolerance (Gallardo and Fereres 1980; Forster et al. 1990), resistance to pests and diseases (Martín et al. 1996) and high seed carotenoid content (Atienza et al. 2004). Several substitution and addition lines have been developed to transfer these traits into wheat (Miller et al. 1982; Fernández and Jouve 1988), and recently, some accessions of *H. chilense* have been used to produce male sterility in wheat with the aim of establishing a wheat hybrid system (Martín et al. 2008; Martín et al. 2010; Castillo et al. 2014). In addition, a major interest in this species relates to its high crossability with other members of the Triticeae tribe (von Bothmer et al. 1986; Martín et al. 1998). Evidence of this is the obtaining of fertile amphiploids named tritordeum (*×Tritordeum* Ascherson et Graebner), which were obtained after chromosome doubling of hybrids between *H. chilense* and tetraploid and hexaploid wheats (Martín and Sánchez-Monge Laguna, 1982; Padilla and Martín 1983; Martín et al. 1999). In addition to *H. chilense*, other *Hordeum* species have been hybridized with wheat; however, the only amphiploid showing enough fertility to be considered as a possible crop is *×Tritordeum martinii* (H^ch^H^ch^AABB) (Pujadas Salvá 2016), which resulted from interspecific crosses between *H. chilense* and cultivated durum wheat. Today, *Tritordeum martinii* is grown in Spain, France, and Italy, being the only synthetic amphiploid crop commercialized for human consumption.

*Hordeum chilense* is an extremely variable species, which can be found in many contrasting environments (Tobes et al. 1995; Giménez et al. 1997. It shows a very wide range of variation of morphological characters (von Bothmer et al. 1980, 1995), different levels of avoidance to barley leaf rust (Rubiales and Niks 1992, 1996) and it even differs at the cytoplasmic level (Atienza et al. 2007; Martín et al. 2009). In 2001, a study concluded that *H. chilense* consists of at least three morphologically and genetically distinct subspecific taxa (Vaz Patto et al. 2001). However, not much work has been done to identify or define further these three subspecific taxa or groups.

*Hordeum chilense*, and the genus *Hordeum* in general, has been characterized by FISH using several repetitive sequences. However, most of the repetitive probes used in this genus were aimed at identifying the 7 pairs of *Hordeum* chromosomes, and more repetitive sequences are required to identify individual chromosome arms or smaller regions introgressed into wheat, which ultimately, is the goal in most introgression breeding programs.

In this work, we use fourteen repetitive sequences, for the identification and comparison by FISH, of all individual chromosomes of *H. chilense*, *H. vulgare* and *T. aestivum*. Furthermore, we increase the knowledge of the karyotype of the different *H. chilense* groups described, by using these probes in the analysis of three *H. chilense* accessions (H1, H16 and H7), each one of them belonging to one of the three different groups.

## Materials and methods

### Plant material

*Hordeum vulgare* cv. Betzes, *Tritordeum martinii* “HT27” and three accessions of *Hordeum chilense* (H1, H16 and H7) were kindly provided by Prof. Antonio Martín from the Instituto de Agricultura Sostenible (CSIC, Spain). *Hordeum chilense* accessions H1, H16 and H7 belong to the group I, II and III respectively, described by Vaz Patto et al. (2001). *Triticum aestivum* cv. Chinese Spring was also used in this work.

### Fluorescent in situ hybridization (FISH)

#### Chromosome preparation for FISH

Seeds of each genotype used in this study were germinated on wet filter paper in the dark for 5 days at 4°C, followed by a period of 24h-48h at 25°C under constant light. Root tips were obtained from these seedlings and occasionally, from adult plants grown in pots in a controlled environmental room (16/8 h, light/dark photoperiod at 20 °C day and 15°C night, with 70% humidity). Preparation of chromosome spreads was done following Kato et al. (2004) and King et al. (2017) with slight modifications. These modifications were: roots were mainly excised from seedlings, although occasionally, adult plants were also used in the obtaining of chromosomes spreads. Also, the percentage of cellulase was increased from 2% to 4%, being the final enzymatic cocktail: 1% pectolyase Y23 and 4% cellulose Onozuka R-10 (Yakult Pharmaceutical, Tokyo) solution in 1x citrate buffer.

#### Development of probes and labelling

Fourteen repetitive sequences were used in this work (Table 1). All of them were amplified by PCR and the primer sequences and PCR conditions are described in Table 1. Three different polymerase enzymes were used: BIOTOOLS™ DNA polymerase (Biotools S.A., Madrid, Spain); KAPA3G plant polymerase (VWR International GmbH, Erlangen, Germany); and MyFi DNA polymerase (Bioline USA, Taunton, MA) (Table 1). PCR was performed in 50 μl reaction mixtures according to the manufacturer´s instructions of each polymerase, and PCR products were resolved on 2% agarose gels in 1xTBE, stained with ethidium bromide and visualized under UV light. The PCR products were purified using a Qiagen kit (Qiagen, Hilden, Germany) following the manufacturer’s instructions. The purified PCR products were labelled with either biotin-16-dUTP or digoxigenin-11-dUTP, using the Biotin-nick translation mix or the DIG-nick translation mix, respectively (Sigma, St. Louis, MO, USA) according to the manufacturer’s instructions. Nick translation was performed in a PCR machine at 15°C for 90 min followed by a final step at 68°C for 10 min. *Hordeum chilense* was also used to re-probe the samples after FISH with the repetitive sequences in order to identify the *H. chilense* chromosomes in the tritordeum line (HT27). *Hordeum chilense* DNA was extracted from leaves of young plants following the CTAB method described in Murray and Thomson (1980).

#### FISH and GISH experiments

The FISH protocol used in this work was a combination of two protocols previously described in Cabrera et al. (2002) and King et al. (2017). The FISH procedure was performed over two days. Unless indicated otherwise, washes were carried out at room temperature. First day: Slides were crosslinked in an UV Cross linker at 0.125 Joules for 30 secs (twice). The hybridization mixture consisted of 50% formamide, 10% dextran sulfate, 2x saline sodium citrate (SSC), 0.125% sodium dodecyl sulfate (SDS) and 0.1 mg of salmon sperm DNA. The concentration of each probe in the hybridization mixture is described in Table 1. The hybridization mixture was denatured for 8 min at 80 °C and cooled on ice for 5 min. A 40 μl-aliquot of the hybridization mixture was added to the cross-linked samples and a cover-slip applied. Slides were incubated in an in-situ PCR thermal cycler (Leica Microsystems™ ThermoBrite™, Leica Biosystems, Wetzlar, Germany) at 80°C for 7 min to denature the chromosomes. The hybridization was carried out overnight (20-24h) at 37 °C in a humid chamber. Second day: slides were washed in 2xSSC twice for 5 min at 37°C, in 1xSSC twice for 5 min and in TNT (0.1 mol/L Trix-HCL, 0.15 mol/L NaCL, 0.05% Tween-20) for 5 min. Slides were then blocked in 50% (w/v) dried skimmed milk in TNT for 20 min at 37°C and washed in TNT for 2 min. For the detection of biotin and digoxigenin labelled probes, slides were incubated at 37°C for 45 min with streptavidin-CY3 (Sigma) and antidigoxigenin-FITC (Sigma) in 1x phosphate buffered saline (PBS), respectively (Table 1). Slides were then washed in TNT for 5 min and dehydrated in 70% and 100% ethanol for 1 min. After counter-staining with 4’, 6-diamidino-2-phenylindol (DAPI) for 5 min, slides were washed in water for 5 min, dehydrated again and mounted in Vectashield (Vector Laboratories, Burlingame, CA, USA). In tritordeum “HT27”, all chromosome spreads were re-hybridized following the reprobing method of Heslop-Harrison et al. (1992), in order to identify the *H. chilense* chromosomes in the wheat background.

#### Image acquisition and chromosome identification

Hybridization signals were examined using a Leica DM5500B microscope equipped with a Hamamatsu ORCA-FLASH4.0 camera and controlled by Leica LAS X software v2.0. Digital images were processed using Adobe Photoshop CS5 (Adobe Systems Incorporated, USA) extended version 12.0 × 64.

The ideogram for H1, H16, H7, *H. vulgare* and *T. aestivum* chromosomes was based on the hybridization patterns of the probes used in this work and the morphology of chromosomes previously described (Cabrera et al. 1995; Pedersen and Langridge 1997; Prieto et al. 2004; Kato 2011; Szakács et al. 2013; Komuro et al. 2013; Tang et al. 2014).

## Results

### Repetitive sequences hybridized to mostly terminal and interstitial regions: pAs1 and pTa-535

The pAs1 probe is a repetitive DNA sequence isolated from *Aegilops tauschii* Coss. (formerly known as *Ae. squarrosa* L.) (Rayburn and Gill 1986). The pTa-535 is a 342-bp tandem repeat isolated from *T. aestivum* L. (Komuro et al. 2013).

Both probes show similar hybridization patterns in all the *Hordeum* analysed in this work, H1, H16, H7 and *H. vulgare*, with all the chromosomes showing signals in both arms (Fig. 1). The three accessions of *H. chilense* H1, H16 and H7 show no major differences in their hybridization patterns (Fig. 1). The lowest and weakest number of hybridization signals per chromosome was observed in *H. vulgare* (Fig. 1). Both pAs1 and pTa-535 predominantly hybridized to the telomeric and subtelomeric regions, with some interstitial and centromeric signals in some chromosomes (Fig. 1). In H1, H16 and H7, chromosomes 1H^ch^and 4H^ch^ exhibited weaker signals than the rest of the chromosomes; while in *H. vulgare*, chromosomes 3H^v^, 4H^v^ and 5H^v^ exhibited the weakest signals (Fig. 1).

**Fig. 1.**
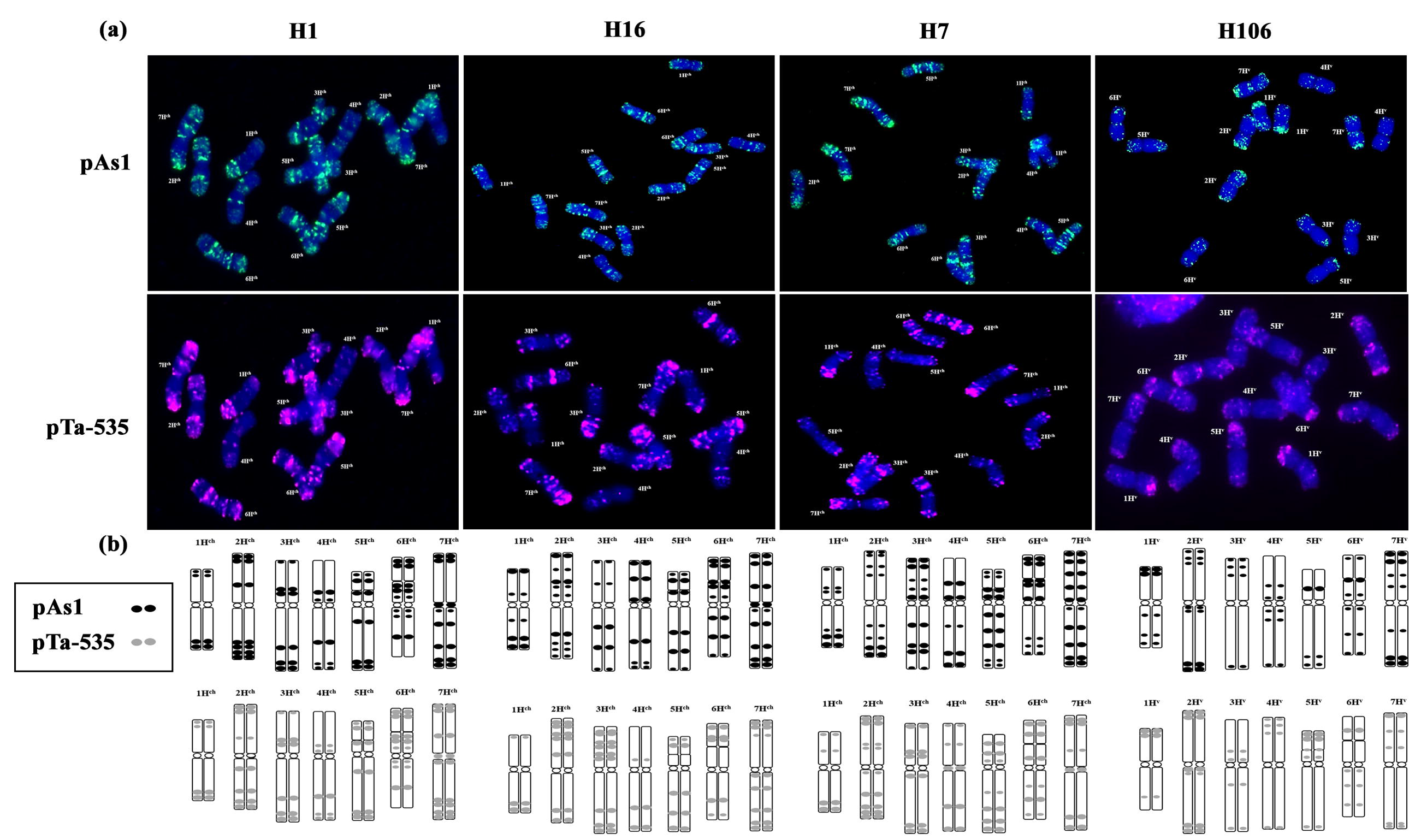
FISH patterns (a) and signal distributions (b) of pAs1 and pTa535 on mitotic metaphase chromosomes of *H. chilense* (H1, H16 and H7) and *H. vulgare*.

In wheat, both probes mainly hybridized to both chromosome arms in all D genome chromosomes as previously described (Rayburn and Gill 1986; Komuro et al. 2013). Signals were also predominantly telomeric and subtelomeric, as was observed in the *Hordeum* analysed. Some chromosomes from the A and B-genomes occasionally hybridized to both probes, but these signals were weak and unsteady, so only D-genome chromosomes were identified in this work.

### Repetitive sequences hybridized to terminal regions: pSc119.2 and HvT01

The pSc119.2 probe is a repetitive sequence containing 120 bp from *Secale cereal* L. (Bedbrook et al. 1980). The HvT01 is a subtelomeric sequence from *H. vulgare* L. (Belostotsky and Ananiev 1990). Hybridization signals with both pSc119.2 and HvT01 were polymorphic in all plant material analyzed in this study (Figs. 2, 5).

**Fig. 2.**
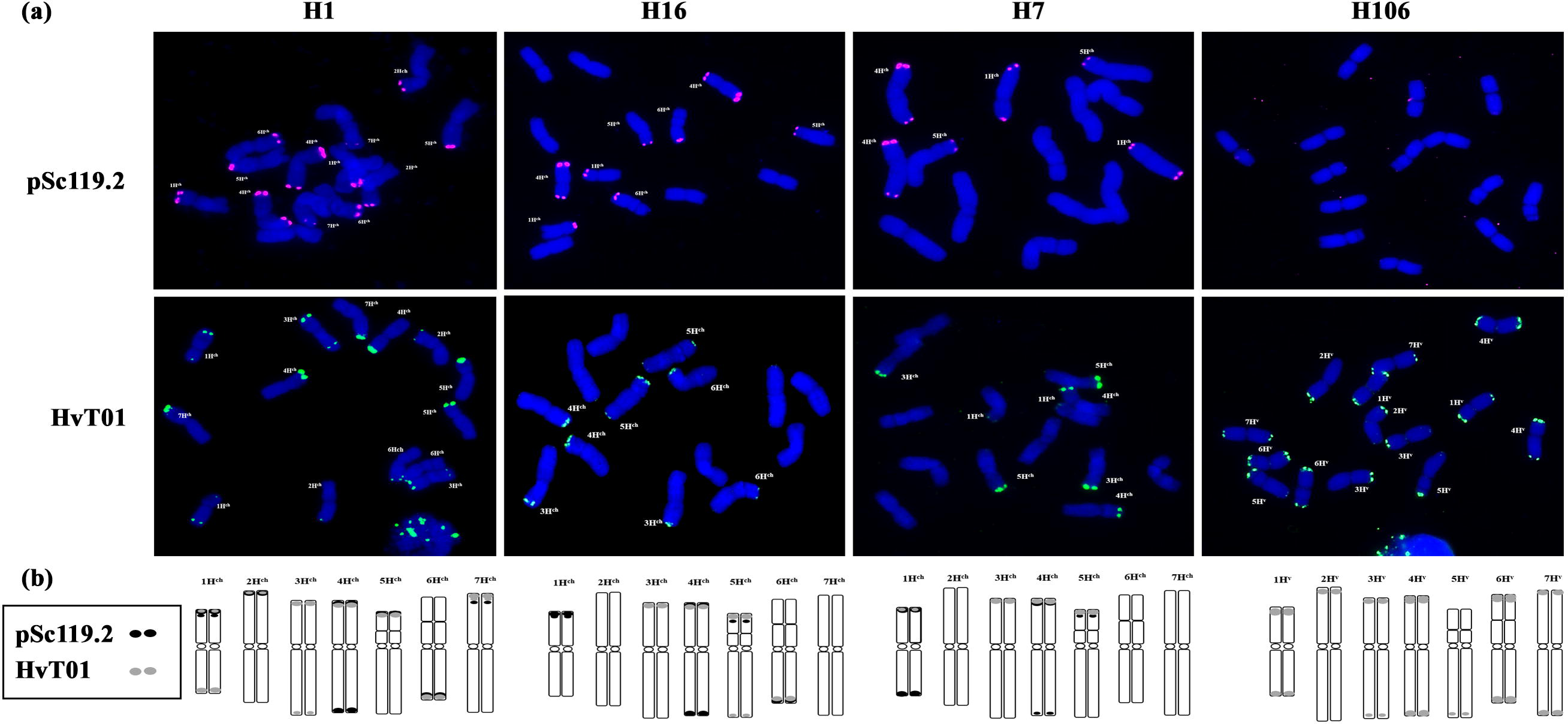
FISH patterns (a) and signal distributions (b) of pSc119.2 and HvT01 on mitotic metaphase chromosomes of *H. chilense* (H1, H16 and H7) and *H. vulgare*.

In *H. chilense* H1, six pairs and seven pairs of chromosomes were hybridized to pSc119.2 and HvT01, respectively (Fig. 2). Probe pSc119.2 did not hybridize on chromosome 3H^ch^, and the signals produced on chromosome 7H^ch^ were weak (Fig. 2). The pSc119.2 signals were detected on the short arm of 1H^ch^, 2H^ch^, 5H^ch^ and 7H^ch^ (1H^ch^S, 2H^ch^S, 5H^ch^S and 7H^ch^S), on the long arm of 6H^ch^ (6H^ch^L) and on both arms of 4H^ch^ (Fig. 2). Probe HvT01 was detected on 2H^ch^S, 4H^ch^S, 5H^ch^S, 6H^ch^L and 7H^ch^S, and on both arms of 1H^ch^ and 3H^ch^ (Fig. 2). In *H. chilense* H16, four pairs of chromosomes were labelled with pSc119.2 and HvT01 (Fig. 2). The pSc119.2 signals were detected on 1H^ch^S, 5H^ch^S, 6H^ch^L and on both arms of 4H^ch^ (Fig. 2). The HvT01 signals were detected on 3H^ch^S, 4H^ch^S, 6H^ch^L and on both arms of 5H^ch^ (Fig. 2). In *H. chilense* H7, both repetitive sequences (pSc119.2 and HvT01) were detected on 3 pairs of chromosomes (Fig. 2). The pSc119.2 signals were observed on 5H^ch^S and on both arms of 1H^ch^ and 4H^ch^ (Fig. 2). The HvT01 signals were detected on 3H^ch^S, 4H^ch^S and 5H^ch^S (Fig. 2). The HvT01 signals obtained in H1 and H7 agreed with the results of Prieto et al. (2004).

In *H. vulgare*, no hybridization signal was detected using the pSc119.2 probe (Fig. 2), which agreed with previous studies (Gupta et al. 1989). However, all pairs of chromosomes were hybridized to HvT01 on both chromosome arms, except 2H^v^S (Fig. 2).

In wheat, thirteen pairs of chromosomes were hybridized to the pSc119.2 (Fig. 5). Signals were detected on all B-genome chromosomes and on chromosomes 4AL, 5AS, 2DS, 3DS, 4DS and 5DS (Fig. 5). All the pSc119.2 signals were detected at the telomeric and subtelomeric regions (Fig. 5) as observed in the genus *Hordeum*. Probe HvT01 did not produce any signal in wheat. Occasionally, two pairs of chromosomes showed a very weak signal, but since the results were not consistent, they were not considered in this work (Fig. 5).

### Repetitive sequences hybridized mostly to interstitial, centromeric and pericentromeric regions: GAA, (AAC)_5_, (CTA)_5_, 4P6 and (AG)_12_

The GAA probe is a GAA-satellite sequence from wheat and barley (Dennis et al. 1980; Pedersen et al. 1996). The (AAC)_5_ probe is a tri-nucleotide repeat from *T. aestivum* L. (Cuadrado et al. 2008). The (CTA)_5_ probe is a tri-nucleotide repeat from *T. aestivum* L., which has been used for the first time in this work. The 4P6 probe is a tandem repeat BAC clone from *Ae. tauschii* Coss. (Zhang et al. 2004). The (AG)_12_ probe is a di-nucleotide repeat from *T. aestivum* L. (Cuadrado et al. 2008).

The GAA sequence was abundant in all the analysed species, allowing for the identification of all *Hordeum* chromosomes and all the wheat B-genome chromosomes (Figs. 3,5). This probe predominantly hybridized to interstitial and pericentromeric regions, with some distal signals in some chromosomes. In *H. chilense* H1, several pericentromeric signals were detected on chromosomes 3H^ch^ and 7H^ch^, and several interstitial signals on 4H^ch^S and 2H^ch^S. Chromosome 1H^ch^L was the only chromosome showing a strong terminal signal. Chromosome 5H^ch^S and 7H^ch^S showed occasionally a weak interstitial and telomeric signal, respectively. In *H. chilense* H16, the hybridization pattern was slightly different to H1. Chromosomes 6H^ch^ and 7H^ch^ showed several pericentromeric and centromeric signals, chromosome 4H^ch^S and 5H^ch^L showed several interstitial signals and chromosome 2H^ch^L showed a strong interstitial signal. Chromosomes 1H^ch^L and 3H^ch^L showed a strong terminal signal. Chromosomes 1H^ch^S and 4H^ch^S showed occasionally a weak telomeric signal, and 2H^ch^L and 7H^ch^L a weak interstitial one (Fig. 3). *Hordeum chilense* H7 showed less signals than H1 and H16. Chromosomes 2H^ch^, 3H^ch^, 4H^ch^, 5H^ch^ and 6H^ch^ showed several pericentromeric signals, and chromosomes 4H^ch^S and 7H^ch^S showed several interstitial signals. Chromosome 2H^ch^L showed a strong interstitial signal. Chromosomes 1H^ch^L, 3H^ch^L and 7H^ch^S showed a terminal signal (Fig. 3). The hybridization pattern detected with the GAA probe on *H. vulgare* was quite different to the pattern observed in *H. chilense*. All chromosomes showed centromeric or pericentromeric signals, with only chromosome 3H^v^L showing a strong distal signal (Fig. 3). As mentioned above, in wheat, the GAA probe hybridized to all B-genome chromosomes, with strong signals distributed along the whole chromosomes (Fig. 5). All chromosomes from the A-genome plus 1DS, 7DS and both arms of 2D, also showed some GAA signal, but the number and intensity were much lower than the ones observed in the B-genome (Fig. 5). The GAA signals pattern agreed with the one described by Pedersen and Langridge (1997).

**Fig. 3.**
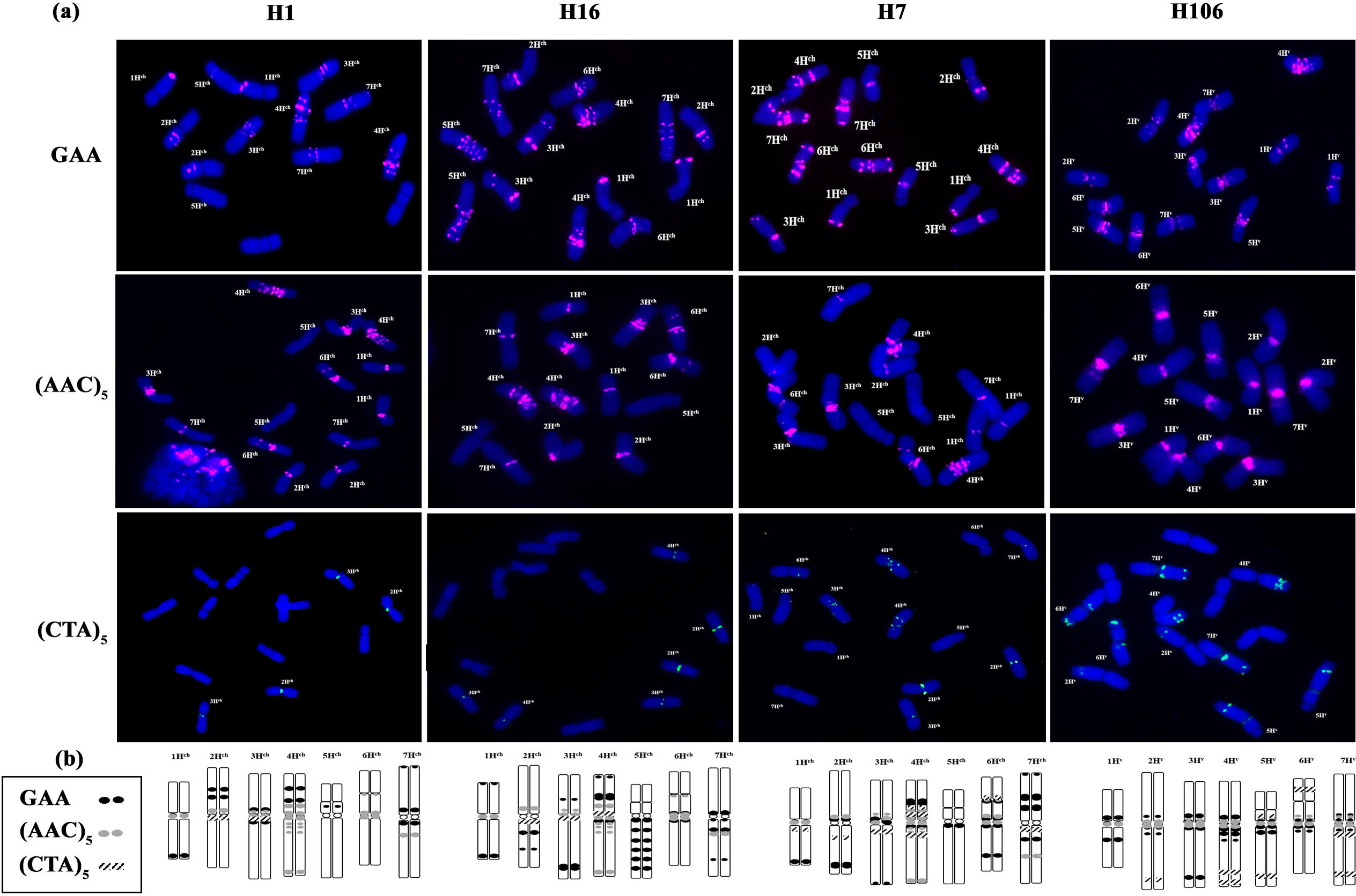
FISH patterns (a) and signal distributions (b) of GAA, (AAC)_5_ and (CTA)_5_ on mitotic metaphase chromosomes of *H. chilense* (H1, H16 and H7) and *H. vulgare*.

The (AAC)_5_ probe produced an intense signal at the centromeric region of all H1, H16 and H7 chromosomes. An exception was chromosomes 5H^ch^, which did not show any signal, and 7H^ch^, which only showed an interstitial signal on the long arm (Fig. 3). This probe was quite conserved in the three *H. chilense* accessions used in this work. The only difference was on chromosome 4H^ch^L, which showed some interstitial signals in H1 and H16 accessions, but not in H7. In *H. vulgare*, the (AAC)_5_ probe hybridized to the centromeric regions of all chromosomes (Fig. 3). In wheat, this probe hybridized to all B-genome chromosomes, and it is mainly distributed around the centromeric region, although some interstitial signals were also observed in some of the chromosomes (Fig. 5). Chromosomes 2AS, 4AL and 7AL also showed (AAC)_5_ signals close to the centromere (Fig. 5).

The hybridization pattern of (CTA)_5_ was similar in H1 and H16, hybridizing both to the centromeric regions of chromosomes 2H^ch^ and 3H^ch^, and to 4H^ch^S in the case of H16 (Fig. 3). However, H7 showed a higher number of signals, with all chromosomes except for 5H^ch^ showing signal (Fig. 3). In H7, signals were pericentromeric on chromosomes 1H^ch^L, 2H^ch^L, 3H^ch^L and 7H^ch^L and on both arms of 4H^ch^. On chromosome 6H^ch^, the signal was located at the NOR region (Fig. 3). *Hordeum vulgare* showed (CTA)_5_ signals on all chromosomes, except for chromosomes 1H^v^ and 3H^v^. It hybridized to the pericentromeric and subtelomeric regions on chromosomes 5H^v^L and 6H^v^S and only to the subtelomeric region on chromosomes 2H^v^L, 4H^v^L and 7H^v^L (Fig. 3). In wheat, the (CTA)_5_ probe hybridized interstitially to chromosomes 2AL, 3AL, 5AL, 7AL and 7BL, and at the centromeric region on 2B and 3B (Fig. 5).

The 4P6 probe did not produce any signal in any of the *Hordeum* used in this study (Fig. S1). This probe only hybridized to the wheat D-genome (Fig. 5). In wheat, signals were detected on five D-genome chromosome pairs: on chromosomes 2DL, 4DL, 5DS, 6DS, and on both arms of chromosome 1D (Fig. 5).

The (AG)_12_ probe, as 4P6, was absent in all *Hordeum* species used in this work (Fig. S1). In wheat, some signals were detected on the pericentromeric region of chromosomes 3BS, 5BS and 6BL (Fig. 5).

### Repetitive sequences hybridized to ribosomal DNA: pTa71 and pTa794

The pTa71 probe is a 9-kb EcoRI fragment of the 18S-25S rDNA isolated from *T. aestivum* (Gerlach and Bedbrook 1979). The pTa794 probe is a 410-bp BamHI fragment of the 5S rDNA isolated from *T. aestivum* (Gerlach and Dyer 1980).

Probe pTa71 did not show any difference among the *Hordeum* genotypes used in this study. Signals were detected on the 2 pairs of chromosomes with nucleolar organizing regions (NOR): 5H^ch^S and 6H^ch^S in *H. chilense*, and 5H^v^S and 6H^v^S in *H. vulgare* (Fig. 4). Our results agreed with the results previously published in numerous cytological studies (Cabrera et al. 1995; Szakács et al. 2013; Delgado et al. 2016).

**Fig. 4.**
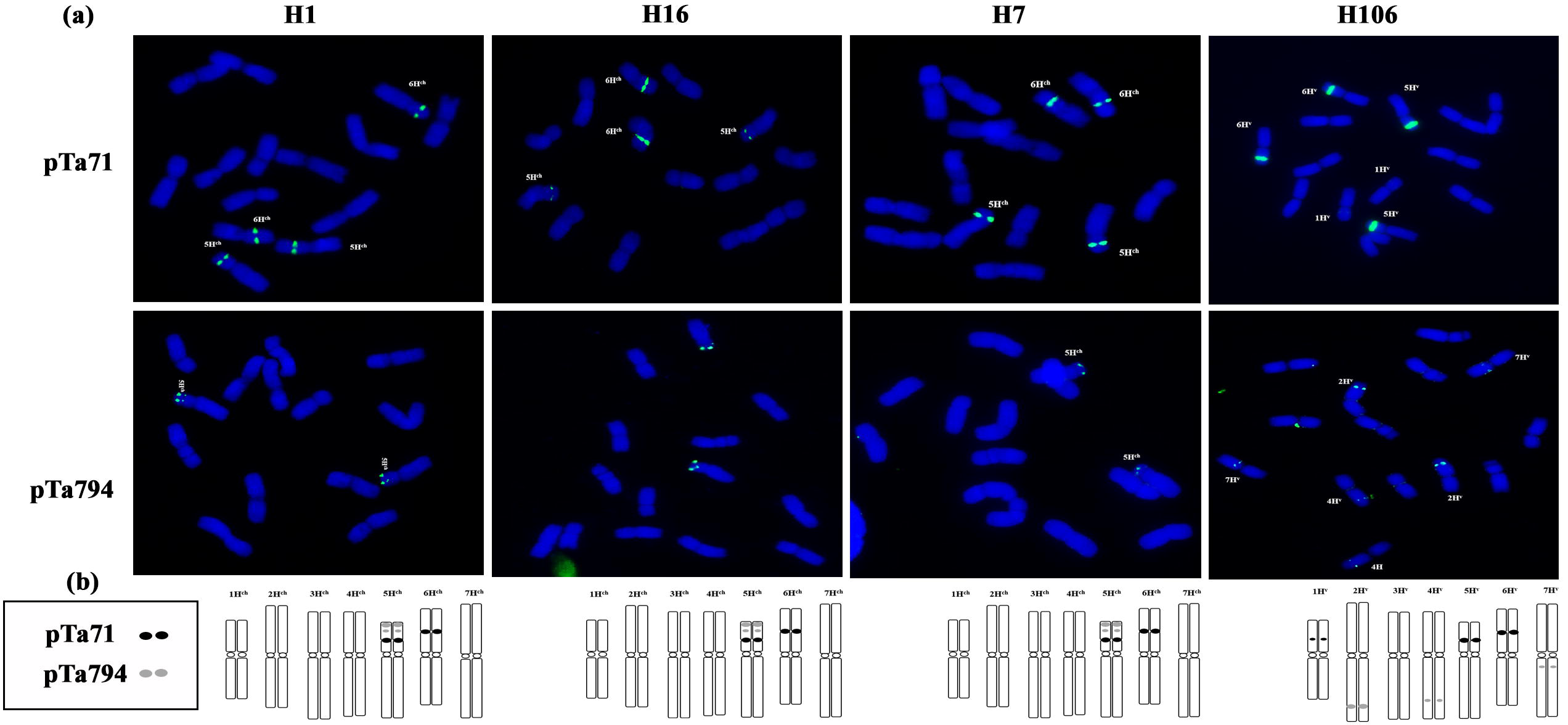
FISH patterns (a) and signal distributions (b) of pTa71 and pTa794 on mitotic metaphase chromosomes of *H. chilense* (H1, H16 and H7) and *H. vulgare*.

On the contrary, the pTa794 probe showed a different pattern among the *Hordeum* studied. In *H. chilense* H1, H16 and H7, this probe was only detected on chromosome 5H^ch^S (Fig. 4). However, in *H. vulgare*, signals were detected on chromosomes 2H^v^L, 4H^v^L and 7H^v^L (Fig. 4). In wheat, the pTa71 probe was detected on chromosomes 1BS, 6BS and 5DS, and the pTa794 probe was observed on chromosomes 1AS, 1BS,1DS, 5AS and 5BS (Fig. 5).

**Fig. 5.**
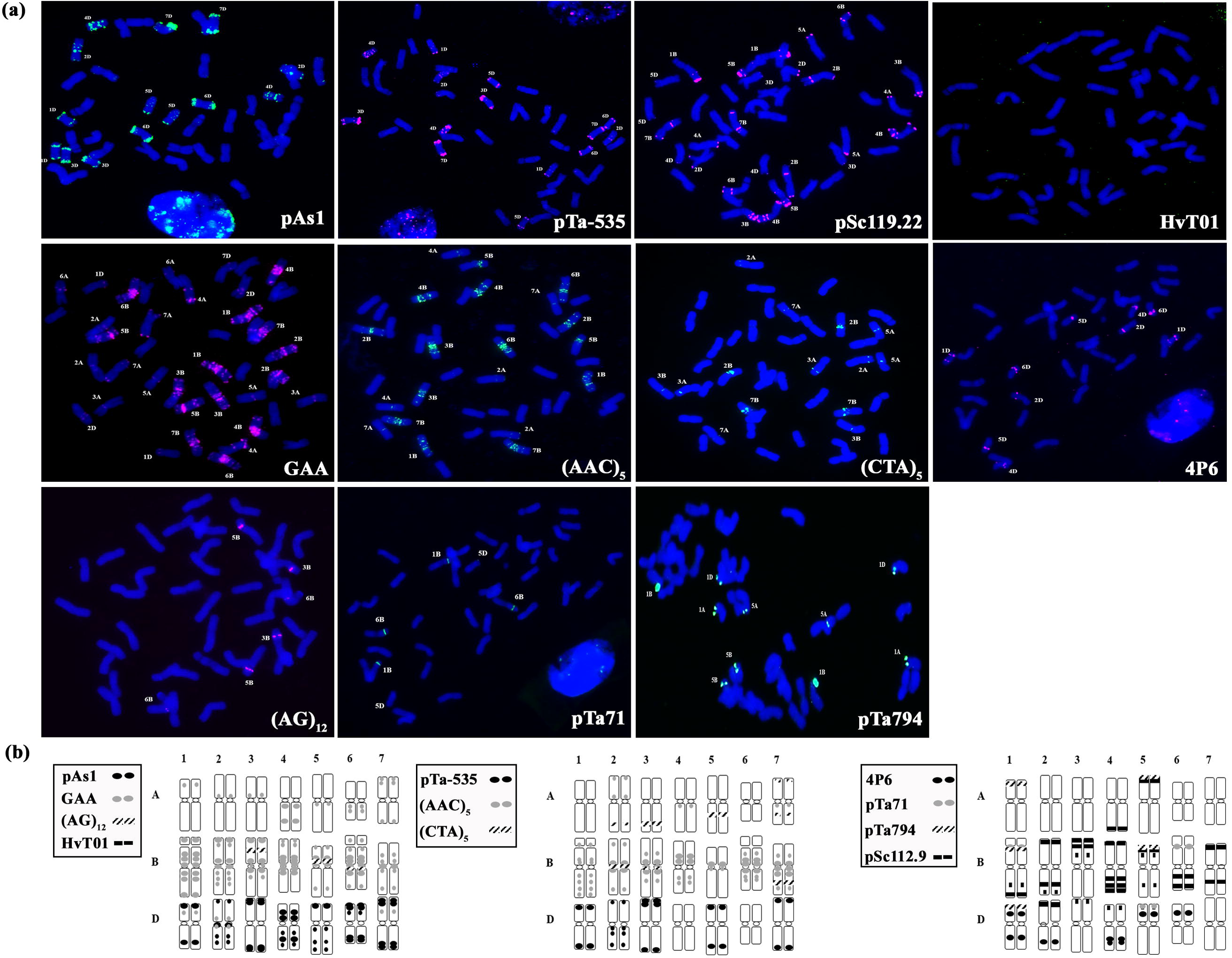
FISH patterns (a) and signal distributions (b) of the fourteen repetitive sequences used in this study on mitotic metaphase chromosomes of *T. aestivum* cv. Chinese Spring.

### Repetitive sequences hybridized to centromeric regions: BAC7, CRW and CCS1

The BAC7 probe is a centromere-specific large insert clone from *H. vulgare* L. (Hudakova et al. 2001). The CRW probe is a wheat centromeric retrotransposon from *Ae. speltoides* Tausch. and *Ae. tauschii* Coss. (Liu et al. 2008). The CCS1 probe is a 260 bp region within the clone (Hi-10) isolated from *B. sylvaticum* L. (Abbo et al. 1995; Aragón-Alcaide et al. 1996).

The centromeric probe BAC7 (Hudakova et al. 2001) was specific to *H. vulgare*, labelling the centromeres of all chromosomes (Fig. 6). Neither *H. chilense* nor *T. aestivum* showed any signals when this probe was used (Fig. 6).

**Fig. 6.**
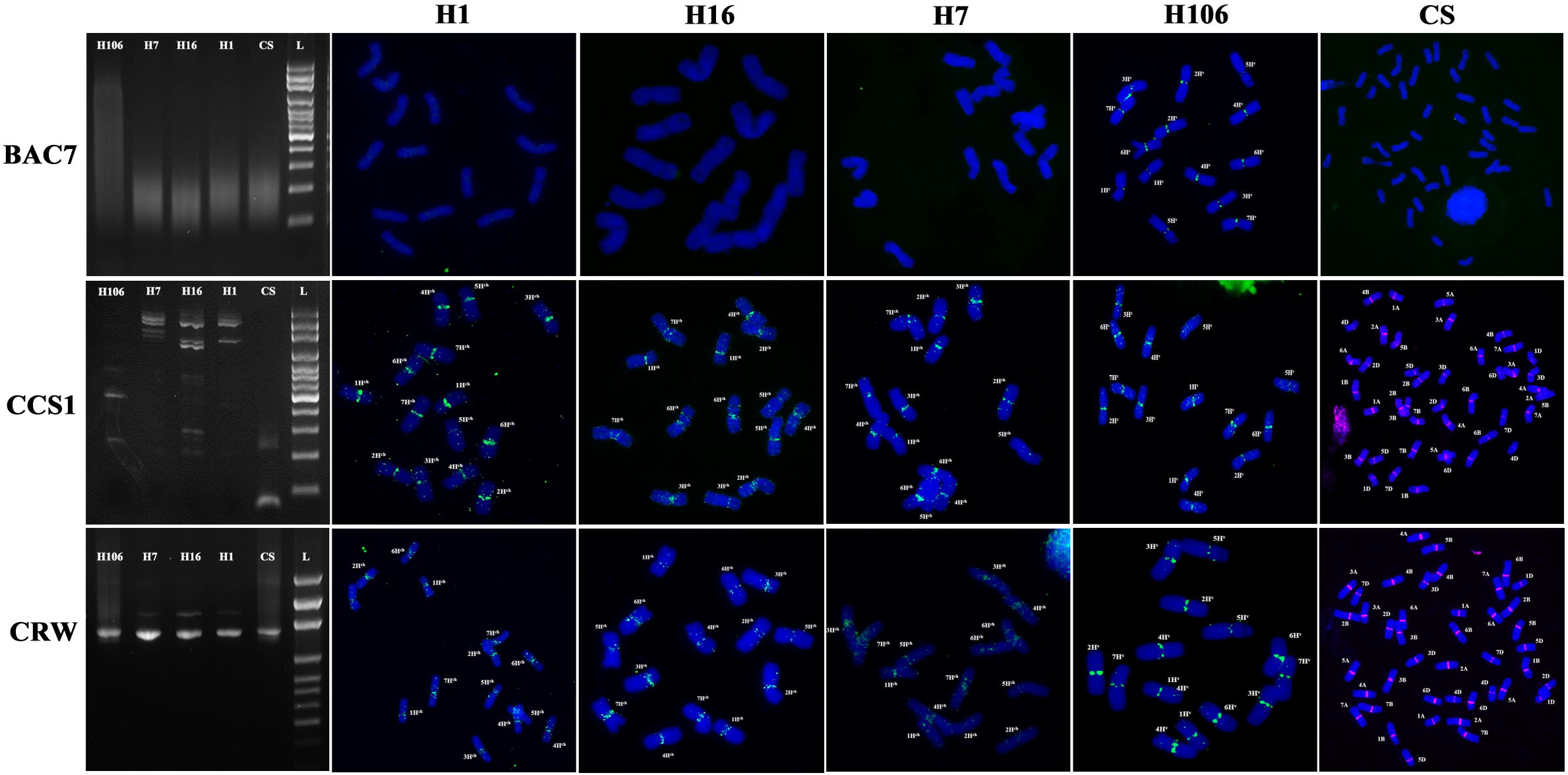
Amplification by PCR and FISH patterns of BAC7, CCS1 and CRW of *H. chilense* (H1, H16 and H7), *H. vulgare* and *T. aestivum* cv. Chinese Spring. The PCR products were visualized on 2 % agarose gels stained with ethidium bromide. L means 100-bp ladder as size marker (Solis BioDyne, Tartu, Estonia).

The centromeric probe CRW (Liu et al. 2008) was detected on all chromosomes of the three accessions of *H. chilense* (H1, H16, H7) and *H. vulgare* (Fig. 6). However, non-specific signals were also frequently observed along chromosomes (Fig. 6). In wheat, probe CRW also hybridized to the centromeric regions of all chromosomes (Fig. 6). Unlike the genus *Hordeum*, signals on wheat were strong and clear, labelling exclusively the centromeric region. Chromosomes from the D-genome showed weaker signals than A and B-genome chromosomes, which is due to the fewer number of CRW copies as previously described (Liu et al. 2008).

The CCS1 probe (Aragón-Alcaide et al. 1996) was also detected in H1, H16, H7, *H. vulgare* and T. aestivum at the centromeric region (Fig. 6). However, as happened with the CRW probe, non-specific signals were frequently observed along chromosomes (Fig. 6). In wheat, the CCS1 pattern was the same as CRW: signals were strong, labelled the centromeric region and chromosomes from the D-genome showed weaker signals than A and B-genome chromosomes (Fig. 6).

Up until now, no *H. chilense* specific centromere probe has been described. Therefore, in an attempt to identify *H. chilense* centromeres in the background of wheat, both probes were tested in the *Tritordeum* line “HT27” (described in Cabo et al. 2014a). All wheat chromosomes showed CRW and CCS1 signals as expected; however, surprisingly, *H. chilense* chromosomes did not show any signal with any of the probes (Fig. S2). On the other hand, the probe (AAC)_5_, which hybridized around the centromeres in these species, showed signals on *H. chilense* chromosomes in the background of bread wheat (Fig. S2).

## Discussion

Triticeae species have large genomes, which are primarily composed of repetitive sequences. Many of these repetitive sequences have been used as probes in FISH analysis for genome differentiation, phylogenetic relationship analysis and chromosome identification among different species of the Triticeae tribe (Cabrera et al. 1995; Taketa et al. 2000; Hagras et al. 2005; Jiang and Gill 2006; Marín et al. 2008; Cuadrado et al. 2013; Komuro et al. 2013). The identification and comparison of *H. chilense* and *H. vulgare* chromosomes have been carried out in several studies (Hagras et al. 2005; Szakács et al. 2013), however, only a small number of probes and one accession of *H. chilense* were used. For the comparison of both *H. chilense* and *H. vulgare* with bread wheat, there is barely any published studies where all individual chromosomes are targeted. In this work, our aim was to provide a clear karyotype of *H. chilense, H. vulgare* and *T. aestivum* chromosomes, which could be useful in wheat breeding programs to monitor *H. chilense* and *H. vulgare* introgressions into wheat, but also to identify the wheat chromosomes where alien segments have been introgressed (Miller et al. 1982; Atienza et al. 2007; Calderón et al. 2012; Rey et al. 2015a, 2015b). Moreover, *H. chilense* and wheat karyotypes would be useful in the development of tritordeum lines, to monitor the chromosome constitution until stable lines are obtained, and to detect spontaneous reorganizations which can occur between *Hordeum* and wheat (Prieto et al. 2001; Cabo et al. 2014b; Delgado et al. 2016; Pujadas Salvá 2016; Delgado et al. 2017).

Here, fourteen repetitive probes were used, which accurately identified all individual chromosomes from three accessions of *H. chilense* (H1, H7 and H16), *H. vulgare* and *T. aestivum*. Briefly, the 4P6 and the (AG_12_) sequences were specific to wheat, and the BAC7 sequence was specific to *H. vulgare*. None of the probes described here was specific to *H. chilense*. The pSc119.2 probe hybridized to both wheat and *H. chilense*, but not to *H. vulgare*. The rest of the probes hybridized to all three species. At the individual chromosome level, using the pSc119.2, the HvT01 and the GAA probes together, we could identify each individual chromosome from the three accessions of *H. chilense* (H1, H16 and H7) and *H. vulgare.* In bread wheat, by using the pAs1 and the GAA probe together we could identify all individual chromosomes, and then by combining them with GISH to label the wheat genome, it was possible to differentiate every wheat chromosome from the ones from barley (both *H. chilense* and *H. vulgare*). The remaining repetitive sequences used in this study add further useful information, which, as indicated before, can be used in phylogenetic relationship analysis and when the identification of small chromosome fragments is required.

Three centromeric probes, BAC7, CCS1 and CRW were used in this study. Probe BAC7 was specific to *H. vulgare* as previously described (Hudakova et al. 2001), so no signal was detected in *H. chilense* and wheat. In the case of both CCS1 and CRW, they are conserved in the three species, labelling the centromeres of all chromosomes (Fig. 6). However, the hybridization signal on wheat was much stronger and clearer than in barley species (H1, H16, H7 and *H. vulgare*), which frequently showed some background signals. Moreover, a confusing result is that neither the CCS1 nor the CRW probes, showed any signal on *H. chilense* chromosomes when present in the background of tritordeum (Fig. S2). In tritordeum line HT27, wheat chromosomes always showed a centromeric signal, however, this signal was absent in *H. chilense* chromosomes. A possible explanation for this result is that barley carries fewer copies of both CCS1 and CRW and when placed in the background of wheat, the signal is too weak to be detected. This is observed in the case of the wheat D-genome, which carries less copies of CCS1 and CRW sequences compared to the A and B-genome, and accordingly, the hybridization signal in weaker (Liu et al. 2008). None of the centromeric sequences described, can therefore be used to identify the centromeric region of *H. chilense* chromosomes in tritordeum lines. However, an alternative is to use the (AAC)_5_ probe. This repetitive sequence, hybridizes to the centromeric and pericentromeric region of all chromosomes of *H. chilense*, except chromosome 5H^ch^ and 7H^ch^, and to all chromosomes of *H. vulgare.* We tested the (AAC)_5_ sequence in tritordeum line HT27, and *H. chilense* chromosomes were perfectly labelled (Fig. S2). Therefore, the (AAC)_5_ sequence can be used to identify the centromeric region of most *H. chilense* chromosomes in tritordeum or any other wheat background.

It has been suggested that *H. chilense* consists of at least three morphologically and genetically distinct subspecific taxa or groups (Vaz Patto et al. 2001). Repetitive DNA sequences are the main components of heterochromatin and are subject to rapid change, therefore, changes in the distribution of repetitive DNA sequences can provide information of genome evolution and speciation. In this study, we selected one accession of each of the three groups described, to check whether the variability observed morphologically and with molecular markers, is also confirmed by FISH using the fourteen repetitive sequences. *Hordeum chilense* H1 belongs to group I, H16 to group II and H7 to group III. The results obtained in this work, support the presence of the three different groups. The centromeric probes and the NOR-associated probes did not show any difference among the different *H. chilense* accessions as expected, since they are very conserved regions. However, the seven repetitive sequences (pAs1, pTa-535, pSc119.2, HvT01, GAA, (AAC)_5_ and (CTA)_5_) did differ. Probe (CTA)_5_ showed the same hybridization pattern in H1 and H16, which differed from the one of H7. In the case of pAs1, pTa-535, pSc119.2, GAA and (CTA)_5_, the hybridization pattern was different in the three *H. chilense* accessions. Based on the FISH results obtained here, accessions H1 and H16 share more similarity between them than with H7. Although these results indicate that the selected set of probes could be useful for a phylogenetic analysis of the different groups described in *H. chilense*, these three accessions are only one of the many accessions included in each of the groups described, and to confirm the existence of different hybridization pattern in each group, more individuals from each group need to be analyzed.

Using the pAs1 probe, Cabrera et al. (1995) suggested that the D-genome from wheat, was the closest phylogenetically to the *H. chilense* genome. We were hoping to add more information supporting this result by using the fourteen probes described in this work. However, only the probes pAs1 and pTa-535, (which show a very similar hybridization pattern to pAs1) have shown some similarity between the D and H^ch^ genomes.

The hybridization pattern obtained in this work agreed with what it has been published before except for the (AG_12_) sequence. We identified three pairs of wheat chromosomes (3BS, 5BS and 6BL) using this probe, while Cuadrado et al. (2008) also describes a signal on chromosome 4B. This difference in the result is probably a consequence of the different FISH conditions used in each study. At this point, it is worth mentioning that some FISH signals can be altered by several factors such as the chromosome spread, the quality of the probe or even the sort of microscope used in the study. Moreover, the examination and identification of the hybridization pattern obtained using repetitive sequences is a complex process that requires experience and previous knowledge of the chromosomal morphology of the species studied. The chromosome spread is critical for obtaining good hybridization patterns when individual chromosomes, chromosome arms or smaller chromosome regions are identified. Some of the hybridization patterns described in this work have already been described previously using different methods for chromosomes spread preparations such as the use of colchicine, ice cold water or nitrous oxide gas (N_2_O) (Cabrera et al. 1995; Taketa et al. 1999, 2000; Komuro et al. 2013; Tang et al. 2016). These three treatments are extensively used in cytogenetic analysis. However, in our experience, N_2_O treatment is the quickest, most reliable and most reproducible method. Here, we combined the protocols from Cabrera et al. (2002) and King et al. (2017) and provide a detailed protocol of how all the FISH experiments were performed in this study.

In summary, we use fourteen repetitive probes to create several karyotypes of *H. chilense* (accessions H1, H16, H7), *H. vulgare* and *T. aestivum;* which together, allow for the identification of every single chromosome in all three species. Moreover, we identify large polymorphism in the three accessions of *H. chilense* studied, which supports the proposal of the existence of different groups inside *H. chilense* species.

## Author contributions

These authors made the following contributions to the manuscript: M-D.R. and A.C.M. designed the research and wrote the manuscript. M-D.R. performed the research and analyzed the data. All authors read and approved the final manuscript.

## Compliance with ethical standards

### Research involving human participants and/or animals

No research involving human participants or animals was performed.

## Acknowledgement

The authors thank Ali Pendle (John Innes Centre, UK) for her valuable comments in the writing of the manuscript, Cai-yun Yang and Dr. Julie King (University of Nottingham (UK)) for allowing to go to their lab to learn the chromosome spread technique using N_2_O treatment and Dr. Angeles Cuadrado (University of Alcalá, (Spain)) for the supply of pTa794 clone. This work was supported by the UK Biotechnology and Biological Research Council (BBSRC) through a grant part of the Designing Future Wheat (DFW) Institute Strategic Programme (<u>BB/P016855/1</u>) and by a Marie Curie Fellowship Grant (H2020-MSCA-IF-2015-703117).

## Tables

**Table 1. List of repetitive sequences with their primer sequences, annealing temperatures and polymerase enzymes used in PCR for the development of FISH probes.** F and R mean forward and reverse primers, respectively.

## Supporting Information

**Fig. S1. Absence of FISH signal using 4P6 and (AG)_12_ probes on mitotic metaphase chromosomes of *H. chilense* (H1, H16 and H7) and *H. vulgare***.

**Fig. S2. FISH pattern of CCS1, CRW and (AAC)_5_ probes and identification by GISH of all *H. chilense* chromosomes on mitotic metaphase chromosomes of tritordeum “HT27”**.

## References

Abbo, S., Dunford, R.E., Foote, T., Reader, S.M., Flavell, R.B., and Moore, G. 1995. Organization of retroelement and stem-loop repeat families in the genomes and nuclei of cereals. Chromosome Res. 3: 5–15.

Aragón-Alcaide, L., Miller, T., Schwatzacher, T., Reader, S., and Moore, G. 1996. A cereal centromeric sequence. Chromosoma, 105: 261–268.

Atienza, S.G., Ávila, C.M., and Martín, A. 2007. The development of a PCR-based marker for PSY1 from Hordeum chilense, a candidate gene for carotenoid content accumulation in tritordeum seeds. Crop. Pasture Sci. 58: 767–773.

Atienza, S.G., Ramírez, C.M., Hernández, P., and Martín, A. 2004. Chromosomal location of genes for carotenoid pigments in Hordeum chilense. Plant Breeding, 123: 303–304.

Bedbrook, J.R., Jones, J., O’Neil, M., Thompson, RD., and Flavell, R.B. 1980. Molecular characterization of telomeric heterochromatin in Secale species. Cell, 19: 545–560.

Belostotsky, D.A., and Ananiev, E.V. 1990. Characterization of relic DNA from barley genome. Theor. Appl. Genet. 80: 374–380.

Cabo, S., Carvalho, A., Martín, A., and Lima-Brito, J. 2014b. Structural rearrangements detected in newly formed hexaploid tritordeum after three sequential FISH experiments with repetitive DNA sequences. J. Genet. 93: 183–188.

Cabo, S., Carvalho, A., Rocha, L., Martín, A., and Lima-Brito, J. 2014a. IRAP, REMAP and ISSR fingerprinting in newly formed hexaploid tritordeum (9Tritordeum Ascherson et Graebner) and respective parental species. Plant Mol. Biol. Rep. 32:761–770.

Cabrera, A., Friebe, B., Jiang, J., and Gill, B.S. 1995. Characterization of Hordeum chilense chromosomes by C-banding and in situ hybridization using highly repeated DNA probes. Genome, 38: 435–442.

Cabrera, A., Martín, A., and Barro, F. 2002. In-situ comparative mapping (ISCM) of Glu-1 loci in Triticum and Hordeum. Chromosome Res. 10(1): 49–54.

Calderón, M.D.C., Ramírez, M.D.C., Martín, A., and Prieto, P. 2012. Development of Hordeum chilense 4Hch introgression lines in durum wheat: a tool for breeders and complex trait analysis. Plant Breeding, 131: 733–738.

Castillo, A., Atienza, S.G., Martín, A.C. 2014. Fertility of CMS wheat is restored by two Rf loci located on a recombined acrocentric chromosome. J. Exp. Bot. 65: 6667–6677.

Cuadrado, A., Cardoso, M., and Jouve, N. 2008. Increasing the physical markers of wheat chromosomes using SSRs as FISH probes. Genome, 51(10): 809–815.

Cuadrado, A., Carmona, A., and Jouve, N. 2013. Chromosomal characterization of the three subgenomes in the polyploids of Hordeum murinum L.: New Insight into the Evolution of This Complex. PLoS ONE, 8(12): e81382.

Delgado, A., Carvalho, A., Martin A.C., Martin, A., and Lima-Brito, J. 2016. Use of the synthetic Oligo‐ pTa535 and Oligo-pAs1 probes for identification of Hordeum chilense-origin chromosomes in hexaploid tritordeum. Genet. Resour. Crop. Evol. 63:945–951.

Delgado, A., Carvalho, A., Martin A.C., Martin, A., and Lima-Brito, J. 2017. Genomic reshuffling in advanced lines of hexaploid tritordeum. Genet. Resour. Crop. Evol. 64(6): 1331–1353.

Dennis, E.S., Gerlach, W.L., and Peacock, W.J. 1980. Identical polypyrimidine-polypurine satellite DNAs in wheat and barley. Hereditas, 44: 349–366.

Fernández, J.A., and Jouve, N. 1988. The addition of Hordeum chilense chromosomes to Triticum turgidum conv. durum. Biochemical, karyological and morphological characterization. Euphytica, 37: 247–259.

Forster, B.P., Phillips, M.S., Miller, T.E., Baird, E., and Powell, W. 1990. Chromosome location of genes controlling tolerance to salt (NaCl) and vigour in Hordeum vulgare and H. chilense. Heredity, 65: 99–107.

Gallardo, M., and Fereres, E. 1989. Drought resistance in tritordeum (Hordeum chilense × Triticum turgidum) in relation to wheat, barley and triticale. Invest. Agrar. Prod. Prot. Veg. 4: 361–37507.

Gerlach, W.L., and Bedbrook, J.R. 1979. Cloning and characterization of ribosomal RNA genes from wheat and barley. Nucleic Acids Res. 7: 1869–1885.

Gerlach, W.L., and Dyer, T.A. 1980. Sequence organization of the repeating units in the nucleus of wheat that contain 5S rRNA genes. Nucleic Acids Res. 8: 4851–4865.

Giménez, M.J., Cosío, F., Martínez, C., Silva, F., Zuleta, A., and Martín, L.M. 1997. Collecting Hordeum chilense Roem et Schult. germplasm in desert and steppe dominions of Chile. Plant Genet. Resour. Newsl. 109: 17–19.

Gupta, S.B. 1969. Duration of mitotic cycle and regulation of DNA replication in Nicotiana plumbaginifolia and a hybrid derivative of N. tabacum showing chromosome instability. Can. J. Genet. Cytol. 11: 133–142.

Hagras, A.A.A., Kishii, M., Tanaka, H., Sato, K., and Tsujimoto, H. 2005. Genomic differentiation of Hordeum chilense from H. vulgare as revealed by repetitive and EST sequences. Genes Gent. Syst. 80: 147–159.

Heslop-Harrison, J.S., Harrison, G.E., and Leitch, I.J. 1992. Reprobing of DNA: DNA in situ hybridization preparations. Trends Genet. 8: 372–373.

Hudakova, S., Michalek, W., Presting, G.G., Hoopen, R.T., Santos K.D., Jasencakova, Z., and Schubert, I. 2001. Sequence organization of barley centromeres. Nucleic Acids Res. 29(24): 5029–5035.

Jiang, J., and Gill, B.S. 2006. Current status and the future of fluorescence in situ hybridization (FISH) in plant genome research. Genome, 49(9): 1057–1068.

Kato, A. 2011. High-density fluorescence in situ hybridization signal detection on barley (Hordeum vulgare L.) chromosomes with improved probe screening and reprobing procedures. Genome, 54: 151–159.

Kato, A., Lamb, J.C. and Birchler, J.A. 2004. Chromosome painting using repetitive DNA sequences as probes for somatic chromosome identification in maize. Proc. Natl Acad. Sci. USA 101: 13554–13559.

King, J., Grewal, S., Yang, C.Y., Hubbart, S., Scholefield, D., Ashling, S., Edwards, K.J., Allen, A.M., Burridge, A., Bloor, C. Davassi A, da Silva, G.J., Chalmers K., and King I.P. 2017. A step change in the transfer of interspecific variation into wheat from Amblyopyrum muticum. Plant Biotech J. 15: 217–226.

Komuro, S., Endo, R., Shikata, K., and Kato, A. 2013. Genomic and chromosomal distribution patterns of various repeated DNA sequences in wheat revealed by a fluorescence in situ hybridization procedure. Genome, 56(3): 131–137.

Liu, Z., Yue, W., Li, D., Wang, R.R.C., Kong, X., Lu, K., Wang, G., Dong, Y., Jin, W., and Zhang, X. 2008. Structure and dynamics of retrotransposons at wheat centromeres and pericentromeres. Chromosoma, 117: 445–456.

Marín, S., Martín, A., and Barro, F. 2008. Comparative FISH mapping of two highly repetitive DNA sequences in Hordeum chilense (Roem. et Schult.). Genome, 51(8): 580–588.

Martín, A., Álvarez, J.B., Martín, L.M., Barro, F., and Ballesteros J. 1999. The development of Tritordeum: A novel cereal for food processing. J. Cereal Sci. 30(2): 85–95.

Martín, A., and Sánchez-Monge Laguna, E. 1982. Cytology and morphology of the amphiploid Hordeum chilense ×Triticum turgidum conv. durum. Euphytica, 31: 261–267.

Martín, A., Martín, L. M., Cabrera, A., Ramírez, M. C., Jimenez, M. J., and Rubiales, D. Hernández, P., and Ballesteros, J. 1998. The potential of Hordeum chilense in breeding Triticeae species. Paper Presented at the Triticeae III (Enfield, NH: Science Publishers), 377–386.

Martín, A., Martínez-Araque, C., Rubiales, D., and Ballesteros, J. 1996. Tritordeum: triticale’s new brother cereal. In: Guedes-Pinto H, Darvey N, Carnide VP (eds) Triticale: Today and Tomorrow, pp. 57–72. Kluwer Academic Publishers, Dordrecht, Netherlands.

Martín, A.C., Atienza, S.G, Ramírez, M.C., and Barro, F. 2008. Male fertility restoration of wheat in Hordeum chilense cytoplasm is associated with 6HchS chromosome addition. Aust. J. Agric. Res. 59: 206–213.

Martín, A.C., Atienza, S.G, Ramírez, M.C., Barro, F., and Martín, A. 2009. Chromosome engineering in wheat to restore male fertility in the msH1 CMS system. Mol. Breed. 24(4): 397–408.

Martín, A.C., Atienza, S.G, Ramírez, M.C., Barro, F., and Martín, A. 2010. Molecular and cytological characterization of an extra acrocentric chromosome that restores male fertility of wheat in the msH1 CMS system. Theor. Appl. Genet. 121(6):1093–101.

Miller, T.E., Reader, S.M., and Chapman, V. 1982. The addition of Hordeum chilense chromosomes to wheat. In: Proc. Int. Symp. Eucarpia on Induced Variability in Plant Breeding (ed. C Broertjes). Pudoc, Wageningen, p. 79–81.

Murray, M.G., and Thompson, W.F. 1980. Rapid isolation of high molecular weight plant DNA. Nucleic Acids Res. 8: 4321–4326.

Padilla, J.A., and Martín, A. 1983. Morphology and cytology of Hordeum chilense x Hordeum bulbosum hybrids. Theor. Appl. Genet. 65(4): 535–355.

Pedersen, C., and Langridge, P. 1997. Identification of the entire chromosome complement of bread wheat by two color FISH. Genome, 40: 589–593.

Pedersen, C., Rasmussen, S.K., and Linde-Laursen, I. 1996. Genome and chromosome identification in cultivated barley and related species of the Triticeae (Poaceae) by in situ hybridization with the GAA-satellite sequence. Genome, 39: 93–104.

Prieto, P., Martín, A., and Cabrera, A. 2004. Chromosomal distribution of telomeric and telomeric associated sequences in Hordeum chilense by in situ hybridization. Hereditas, 141: 122–127.

Prieto, P., Ramírez M.C., Ballesteros, J., Cabrera, A. 2001. Identification of intergenomic translocations involving wheat, Hordeum vulgare and Hordeum chilense chromosomes by FISH. Hereditas, 135(2-3): 171–174.

Pujadas Salvá, A.J. 2016. Notulae taxinomicae, Chorologicae, Nomenclaturales, Bibliographicae aut philogicae opus “Flora Ibérica” intendentes (44-46). 44. × TRITORDEUM MARTINII A. PUJADAS (POACEAE) NOTHOSP. NOV. Acta Bot. Malac. 41: 325–338.

Rayburn, A.L., and Gill, B.S. 1986. Molecular identification of the D-genome chromosomes of wheat. J. Hered. 77: 253–255.

Rey, M.D., Calderón, M.C., and Prieto, P. 2015a. The use of the ph1b mutant to induce recombination between the chromosomes of wheat and barley. Front. Plant. Sci. 6: 160.

Rey, M.D., Calderón, M.C., Rodrigo, M.J., Zacarías, L., Alós, E., and Prieto, P. 2015b. Novel bread wheat lines enriched in carotenoids carrying Hordeum chilense chromosome arms in the ph1b background. PLoS ONE, 10(8); e0134598.

Rubiales, D., and Niks, R.E. 1992. Histological responses in Hordeum chilense to brown and yellow rust fungi. Plant Pathol. 41(5): 611–617.

Rubiales, D., and Niks, R.E. 1996. Avoidance of rust infection by some genotypes of Hordeum chilense due to their relative inability to induce the formation of appressoria. Physiol. Mol. Plant Pathol. 49(2): 89–101.

Szakács, É., Kruppa, K. and Molnár-Láng, M. 2013. Analysis of chromosomal polymorphism in barley (Hordeum vulgare L. ssp. vulgare) and between H. vulgare and H. chilense using three-color fluorescence in situ hybridization (FISH). J. Appl. Genetics 54: 427–433.

Taketa, S., Ando, H., Takeda, K., Harrison, G. E. and Heslop-Harrison, J. S. 2000. The distribution, organization, and evolution of two abundant and widespread repetitive DNA sequences in the genus Hordeum. Theor. Appl. Genet. 100:169–176.

Taketa, S., Harrison, G. E., and Heslop-Harrison, J. S. 1999. Comparative physical mapping of the 5S and 18S-25S rDNA in nine wild Hordeum species and cytotypes. Theor. Appl. Genet. 98: 1–9.

Tang, S., Qiu, L., Xiao, Z., Fu, Su., and Tang Z. 2016. New oligonucleotide probes for ND-FISH analysis to identify barley chromosomes and to investigate polymorphisms of wheat chromosomes. Genes, 7: 118. doi:10.3390/genes7120118.

Tang, T., Yang, Z., and Fu, S. 2014. Oligonucleotides replacing the roles of repetitive sequences pAs1, pSc119.2, pTa-535, pTa71, CCS1, and pAWRC.1 for FISH analysis. J. Appl. Genetics 55: 313–318.

Tobes, N., Ballesteros, J., Martínez, C., Lovazzano, G., Contreras, D., Cosio, F., Gastó, J., and Martín, L.M. 1995. Collection mission of Hordeum chilense Roem. et Schult. in Chile and Argentina. Report of a collecting mission. Genet. Resour. Crop. Evol. 42(3): 211–216.

Vaz Patto, M.C., Aardse, A., Buntjer, J., Rubiales, D., Martín, A., and Niks, R.E. 2001. Morphology and AFLP markers suggest three Hordeum chilense ecotypes that differ in avoidance to rust fungi. Can. J. Bot. 79: 204–213.

Von Bothmer, R., Jacobsen, N., and Jorgensen, R. B. 1986. Taxonomy, variation, and relationships in the Hordeum parodii group (Poaceae). Nord. J. Bot. 6: 399–410.

Von Bothmer, R., Jacobsen, N., and Nicora, E. 1980. Revision of Hordeum sect. Anisolepis Nevski. Bot. Not. 133: 539–554.

von Bothmer, R., Jacobsen, N., Baden, C., Jorgensen, R.B., and Linde-Laursen, I. 1995. An ecogeographical study of the genus Hordeum, 2nd edn. Systematic and ecogeographic studies on crop genepools 7, Rome. ISBN-13: 978-932-9043-229-6.

Zhang, P., Li, W., Fellers, J., Friebe, B., and Gill, B.S. 2004. BAC-FISH in wheat identifies chromosome landmarks consisting of different types of transposable elements. Chromosoma, 112: 288–299.

